# A Portable RPA-CRISPR/Cas12a-Based Biosensing Platform for On-Site Detection of the Microcystin Synthetase E Gene in Lake Water

**DOI:** 10.1101/2025.07.11.664500

**Authors:** Rakibul Hasan, MacKenzie Smith, Siwen Wang

## Abstract

Microcystins are potent cyanotoxins produced by toxigenic cyanobacteria during harmful algal blooms (HABs), posing risks to ecosystems and human health. In this study, we developed a portable RPA-CRISPR/Cas12a biosensing platform for the rapid, on-site detection of the microcystin synthetase E (*mcyE*) gene, a key biomarker for microcystin-producing strains. The developed RPA-CRISPR/Cas12a assays enable detection of the *mcyE* gene within 50 min, with either fluorescence or lateral flow assay readouts. The fluorescence readouts have an analytical detection limit of 48.4 copies/µL and a dynamic range of 1.2 × 10^2^ to 1.2 × 10^7^ copies/µL. To enable field deployment, a magnetic bead-based DNA extraction method was integrated, achieving extraction within 1 hour without centrifugation. The complete workflow demonstrated a method LOD of 8.4 × 10^2^ cells/mL in spiked lake water. Applicability was validated using non-spiked environmental water samples collected from multiple HAB-affected lakes. Importantly, a systematic matrix effect assessment was conducted for the CRISPR sensing step, evaluating environmental variables such as pH, ions, nutrients, and natural organic matter. This study establishes a practical, sensitive, and selective detection tool for proactive HAB monitoring. The platform’s simplicity, portability, and completeness, from sample pretreatment to signal readout, highlight its potential for real-world environmental biosensing applications.

## 1. Introduction

Over the past several decades, harmful algal blooms (HABs) caused by cyanobacteria have become increasingly common and severe in freshwater lakes worldwide, producing cyanotoxins that pose serious threats to both the environment and public health.^1,2^ In the United States alone, the economic burden of HABs is estimated at approximately $100 million annually, taking into account costs associated with public health, commercial fisheries, recreation and tourism, and monitoring and mitigation efforts. ^3^ Among cyanotoxins, microcystins (MCs) are the most prevalent and are produced by various genera of filamentous cyanobacteria. ^4^ While MCs are primarily hepatotoxins, they can also accumulate in other organs and tissues. ^4,5^ Human exposure mainly occurs through drinking water and recreational activities in contaminated freshwater bodies. ^5,6^ Due to their widespread occurrence and significant toxicity, the World Health Organization has set a guideline limit of 1 µg/L for MCs in drinking water. ^5^

Current MC monitoring strategies primarily focus on direct toxin analysis, which only indicates toxin presence once an HAB event has occurred. This approach cannot serve as an early warning system, since MC production is cell-density dependent and typically triggered during peak cell proliferation. ^7^ Early detection of toxin-producing cyanobacteria, before or at the early stages of bloom formation, could provide critical predictive value. Because toxigenic and non-toxigenic strains are often morphologically indistinguishable, genetic identification via microcystin synthetase (*mcy*) genes is a common and effective method. ^8,9^ Among different *mcy* genes (*mcyA* to *J*) ^8^, *mcyE* is a well-proven strong candidate for monitoring due to its essential role in incorporating toxicity-determining amino acids, Adda and D-Glu, into the MC structure, though environmental factors may influence this expression. ^6, 8, 13^ At present, *mcy* gene detection still relies on conventional polymerase chain reaction (PCR)-based methods with thermal cycling for amplification followed by a fluorescent detection system, *e.g.*, real-time quantitative PCR (qPCR), which is the gold standard method for nucleic acid amplification and detection. However, it relies on thermal cycling with an extended reaction time of 1-2 hours, which limits its application for field use or resource-limited settings. ^10^

Biosensing technologies offer promising alternatives that can overcome these limitations and could be a promising supplement to PCR-based methods. Particularly, clustered regularly interspaced short palindromic repeats (CRISPR)/CRISPR-associated nuclease (Cas) sensing systems, originally derived from the adaptive immune system of prokaryotes, ^11,12^ have gained attention for their potential in environmental monitoring. ^13,14^ Platforms such as SHERLOCK (Cas13a) ^15^ and DETECTR (Cas12a) ^16^ incorporate a pre-amplification strategy of recombinase polymerase amplification (RPA), which is favored for its near-ambient reaction temperature, rapid reaction time, and lyophilization compatibility. CRISPR-Cas biosensors utilize a Cas protein and a guide RNA (gRNA), which includes a programmable CRISPR RNA (crRNA) to ensure target specificity, and a structural component that stabilizes the Cas-gRNA complex. ^12^ Upon recognition of a target sequence by the crRNA, the Cas protein is activated, enabling cis-cleavage of the target and non-specific trans-cleavage of nearby single-stranded DNA (ssDNA) or RNA (ssRNA) (*e.g.*, ssDNA for Cas12a and Cas14, and ssRNA for Cas13), which underpins signal generation and amplification. ^12^ Despite significant advances in CRISPR-sensing over the past decade, ^17–22^ its environmental application remains nascent, and it has not been explored for any specific cyanotoxin-producing genes. Substantial challenges arise from the complex nature of environmental water samples, which are characterized by low and fluctuating target concentrations and diverse matrix components. To date, no study has systematically addressed the matrix effects in real environmental water for CRISPR-based detection. Additionally, efficient sample pretreatment, especially the lysis of robust cyanobacterial cells, remains a critical bottleneck for field implementation.

In this study, we developed a fully integrated, portable RPA/CRISPR-Cas12a biosensing platform with dual-mode readouts, i.e., fluorescence and lateral flow assay (LFA), for the on-site *mcyE* gene detection in real lake water samples. The RPA/CRISPR-Cas12a detection assay was developed and optimized using DNA extracted from *Microcystis aeruginosa* PCC 7806 (*M. aeruginosa* PCC 7806). To better understand the matrix effects on CRISPR-sensing, potential influential matrix factors, including pH, alkalinity, natural organic matter (NOM), ammonia nitrogen, phosphorus, Ca^2+^, and Fe^3+^, with environmentally relevant ranges in freswater, were comprehensively investigated. ^23,24^ Furthermore, we developed a magnetic bead-based on-site DNA extraction method and demonstrated the developed CRISPR-based detection platform using both spiked and non-spiked real lake water.

## 2. Materials and Methods

### 2.1. Materials and Reagents

*M. aeruginosa* PCC 7806 was obtained from the Institut Pasteur, France. BG-11 medium was purchased from Sigma-Aldrich (MA, USA). PrimeSTAR HS (Premix) was purchased from TAKARA Bio Inc. The RPA reactions were performed using the TwistAmp™ Basic Kit (TwistDx™, Cambridge, UK). LFA strips (HybriDetect-Universal Lateral Flow Assay Kit) were obtained from Milenia Biotec (Giessen, Germany). All primers, probes, and crRNAs were designed and purchased from Integrated DNA Technologies IDT (IA, USA). NEBuffer 2.1 (10×) and LbCas12a were purchased from New England Biolabs (MA, USA). The GoTaq® Probe qPCR master mix was purchased from Promega (WI, USA). PCR purification kits and QIAamp DNA Mini Kits were purchased from Qiagen (MD, USA). The Dynabeads® DNA DIRECT™ Universal Kit was purchased from Invitrogen by Thermo Fisher Scientific (MA, USA). Sodium dodecyl sulfate (SDS), 0.40 µm polycarbonate membranes (47 mm diameter), and PES membrane filters (0.45 µm pore size, 13 mm diameter, hydrophilic) were sourced from Millipore Sigma (MA, USA). The standard natural organic matter (NOM), Suwannee River NOM, was purchased from the International Society for Humic Substances. All other chemicals were purchased from Sigma Aldrich (MO, USA) and Fisher Scientific (MA, USA).

### 2.2. Sample Collection

Lake water samples were collected from Lake Neatahwanta, Lake Hyde, and Lake Ontario. The coordinates of the sampling locations with the physicochemical parameters of the water samples, including pH, ambient and water temperatures, phosphorus, and nitrogen concentrations, are shown in **Table S1**. Samples were stored at 4 °C in plastic bottles until further analysis.

### 2.3 Cyanobacterial cultivation and DNA extraction

*M. aeruginosa* PCC 7806 was cultivated in BG-11 medium in a shaker incubator (Innova S44i, Eppendorf) under a 12 h light/12 h dark photoperiod at 22 °C with a photon flux density of 5 µmol photons m^-2^ s^-1^. BG-11 medium was refreshed biweekly, and cultures were manually agitated once a week. Genomic DNA was extracted from *M. aeruginosa* PCC 7806 using the commercial kit, i.e., QIAamp DNA Mini Kit. In this DNA extraction method, algal cells were harvested by centrifuging the culture at 10,000×g for 10 min, and the resulting pellet was resuspended in 180 µL of lysozyme solution with an incubation at 37 °C for 30 min and then extracted following the manufacturer’s protocol. The DNA purity and concentration were assessed using a NanoDrop One^C^ (Thermo Scientific, MA, USA).

### 2.4. Design and optimization of crRNA and target sequence

Five crRNAs, crRNA 1-5 (**Table S2**), targeting distinct regions of the *mcyE* gene were designed using the CHOPCHOP tool (http://chopchop.cbu.uib.no/). PCR primers were designed using Amplify4 to amplify template dsDNA (∼1000bp) from the extracted genomic DNA of *M. aeruginosa* PCC 7806for each crRNA target sequence, then validated and optimized with agarose gel electrophoresis and sequencing (Azenta Life Sciences, MA, USA), as shown in **Figure S1,** with the results for crRNA 1 as an example. The optimized primer sequences for crRNA 1 is shown in **Table S3**. PCR amplification was conducted in a 50 µL reaction mixture containing 1 µL of forward primer (10 µM), 1 µL of reverse primer (10 µM), 25 µL of PrimeSTAR HS Premix, 22 µL of nuclease-free water, and 1 µL of extracted DNA. Thermal cycling was performed using PXE 0.2 Thermal Cycler (Thermo Fisher Scientific, MA, USA) under the following conditions: 98 °C for 10 s, 55 °C for 5 s, and 72 °C for 1 min/kb, over 30 cycles.

Each crRNA was initially validated via an *in vitro* cleavage assay, followed by agarose gel electrophoresis. The cleavage reaction was prepared using 1 µL of template dsDNA, 1 µL of EnLbCas12a (1 µM), 1 µL of crRNA (1 µM), 1 µL of 10× NEBuffer 2.1, and 6 µL of nuclease-free water. The mixture was incubated at 37 °C for 30 min. The sensitivity of each crRNA was evaluated based on fluorescence intensity with the CRISPR-Cas12a reporter assay that contains a 25 µL reaction mixture of 16 µL of nuclease-free water, 2.5 µL of 10× NEBuffer 2.1, 2 µL of fluorophore-quencher-labeled single-strand DNA (FQ-ssDNA) reporter (5’-FAM-TTATT-BHQ1-3’, 2 µM), 1 µL of LbCas12a (1 µM), 1 µL of crRNA (1 µM), and 2.5 µL of template dsDNA, following the same incubation conditions as above. The fluorescence intensity was measured using an Infinite M Nano^+^ microplate reader (Tecan, NC, USA) with the excitation and emission wavelengths of 485 nm and 535 nm, respectively. The specificity of the optimal Cas12a/crRNA complex was evaluated by assessing its cleavage activity with agarose gel electrophoresis and fluorescence detection against potential interfering genomic targets, including the *mcyA*, *mcyD*, and *cpcB* genes. All the gel electrophoresis experiments mentioned above were carried out with 1.5% TBE agarose gels at 100 V for 60 min using the MyGel InstaView Mini Electrophoresis System (Accuris Instruments by Benchmark Scientific, NJ, USA).

### 2.5. Development of RPA-CRISPR/Cas12a detection platform

RPA reaction was performed according to the manufacturer’s protocol. The reaction mixture consisted of 2.4 µL each of forward and reverse primers (10 µM) designed using Amplify4 (**Table S3**), 29.5 µL of rehydration buffer, 11.2 µL of nuclease-free water, and 2 µL of the extracted genomic DNA. This 47.5 µL mixture was added to the enzyme pellet and mixed thoroughly. Subsequently, 2.5 µL of 280 mM magnesium acetate (MgOAc) was added, bringing the total volume to 50 µL. The RPA reaction was incubated at 37 °C with reaction time optimization of 10 to 20 min. Following the RPA amplification, the CRISPR/Cas12a-based detection with fluorescence readout was conducted under the same conditions as above for the reporter assay, except that the RPA-amplified product was used as template DNA. The CRISPR reaction time was optimized with 15-30 min. For LFA readout, the ssDNA reporter was labeled with FAM at the 5′ end and biotin at the 3′ end (5′-56-FAM-TTATT-biotin-3′). The sensitivity of the constructed RPA-CRISPR/Cas12a detection platform was assessed with a 10-fold serial dilution of the extracted genomic DNA (ranging from 1.2 × 10^0^ to 1.2 × 10^7^ copies/µL) for both fluorescence- and LFA-based formats. The specificity was confirmed by comparing both fluorescence and LFA detection of *mcyE* with that of *mcyA*, *mcyD*, and *cpcB* genes.

### 2.6. Effect of environmental matrix on CRISPR-based detection

The matrix effects that are commonly present in natural freshwater were investigated for CRISPR/Cas12a-based detection with influential parameters, including pH, alkalinity, NOM, Ca^2+^, Fe^3+^, phosphorus, and ammonia nitrogen. Each factor was adjusted or introduced using appropriate chemical surrogates. Different pH conditions were adjusted across the range of 5 to 9 using controlled additions of NaOH and HCl. Alkalinity was modulated using NaHCO_3_ in concentrations ranging from 10 to 400 mg/L as CaCO_3_. NOM was introduced using standard aquatic NOM at concentrations of 1.0 to 30 mg/L. Ca^2+^ and Fe^3+^ were evaluated with CaCl_2_ and FeCl_3_ at the ranges of 1.0-50 mg/L as Ca^2+^ and 0.1-10 mg/L as Fe^3+^, respectively. Ammonia nitrogen was simulated using NH_4_Cl within the range of 0.5 to 25 mg/L as N. Total phosphorus was represented by KH_2_PO_4_, tested from 0.1 to 3.5 mg/L as PO_4_^3-^. All the above stock solutions were prepared with DI water, sterilized with autoclave, and then spiked with PCR-amplified template dsDNA to a concentration of 3 × 10^9^ copies/µL. All the prepared samples with the matrix-free positive control (template dsDNA diluted with nuclease-free water) at the same concentration were tested with the CRISPR/Cas12a assay at 37 °C for 30 min followed by fluorescence detection.

### 2.7. On-site DNA extraction method development

Genomic DNA from *M. aeruginosa* PCC 7806 was extracted using the commercial kit, i.e., Qiagen DNA Extraction mini kit, and a magnetic bead-based on-site DNA extraction method developed in this study in parallel. The on-site DNA extraction method was developed using a modified protocol based on Dynabeads® DNA DIRECT™ Universal Kit. Briefly, 3 mL of lake water was filtered using a syringe-fitted 0.45 µm PES membrane (13 mm in diameter) housed in a polycarbonate syringe holder (Fisher Scientific, MA, USA). The membrane with retained cells was transferred into a sterile 1.5 mL microcentrifuge tube. For cell lysis, 171 µL of nuclease-free water, 9 µL of proteinase K, and 20 µL of 10% SDS were added, followed by incubation at 65 °C for 30 min. Then, 200 µL of resuspended Dynabeads® lysis buffer was added with rapid pipetting and incubated at 65 °C for 15 min. After cooling to room temperature, the Dynabeads-DNA complex was transferred to a fresh tube and placed on a magnetic rack for 2 min. The supernatant was discarded, and the beads were washed three times with 200 µL of 1× washing buffer. Finally, the DNA-bound beads were resuspended in 50 µL of elution buffer and mixed thoroughly by pipetting multiple times. The suspension was incubated at 65°C for 5 min, then magnetically separated, and the supernatant containing the purified DNA was transferred to a new sterile tube. This separation step was repeated twice to maximize DNA recovery. The developed on-site DNA extraction method was validated using both spiked and non-spiked lake water with a comparison of the commercial kit, i.e., QIAamp DNA Mini Kit. Non-spiked water samples were collected from Hyde Lake, Lake Neatahwanta, and Lake Ontario. The spiked water samples were prepared by a 10-fold serial dilution of *M. aeruginosa* PCC 7806 cell culture with lake water that was filtered by a 0.40 µm polycarbonate membrane (47 mm of diameter), to a concentration range of 4.7 × 10^3^ to 4.7 × 10^7^ cells/mL. The extracted DNA was detected by qPCR (MasterCycler RealPlex 4, Eppendorf, MA, USA) with a 20 µL reaction mixture containing 1 µL of forward primer (10 µM), reverse primer (10 µM), and probe (5 µM) mixture, 10 µL of GoTaq® probe qPCR master mix, 7 µL of nuclease-free water, and 2 µL of extracted DNA. The sequences of qPCR primers and probe are listed in **Table S3**. The thermal cycling was performed by 40 cycles of 95 °C for 2 min, 95 °C for 15 s, and 60 °C for 1 min. The cycle number (C_t_) values of no-template controls (NTC) were all >35.

### 2.8. Demonstration on real lake water

The integrated *mcyE* biosensing platform, featuring the developed on-site DNA extraction method and RPA-CRISPR/Cas12a detection, was demonstrated using both spiked and non-spiked lake water samples prepared according to the same method mentioned above. The samples were extracted with the on-site DNA extraction protocol, followed by the RPA-CRISPR/Cas12a biosensing with fluorescence and LFA readouts.

## 3. Results and discussion

### 3.1. Mechanism and workflow of the RPA-CRISPR/Cas12-based mcyE gene detection

The detection mechanism of the RPA-CRISPR/Cas12a biosensor for the *mcyE* gene is illustrated in **Figure 1a**. Following sample collection, total DNA is extracted using a magnetic bead-based protocol (detailed in **Figure 1b**) developed in this study to efficiently lyse cyanobacterial cells and recover nucleic acids under field-compatible conditions. The validation, optimization, and performance of the RPA-CRISPR/Cas12a biosensing assay and the on-site DNA extraction method will be further discussed in the following sections.

**Figure 1.**
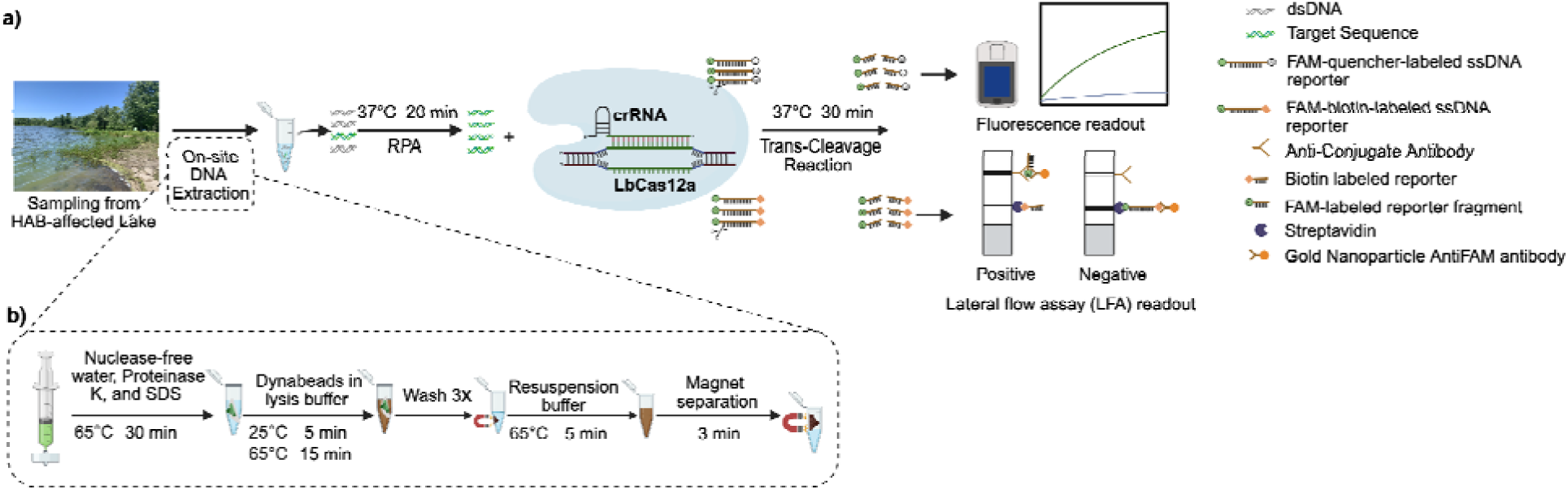
Schematic illustration of the on-site RPA-CRISPR/Cas12a biosensing platform for the *mcyE* gene detection in lake water. a) RPA/CRISPR-Cas12a-based biosensing diagram with dual-mode readouts through fluorescence and lateral flow assay (LFA). b) On-site DNA extraction method for cyanobacteria from lake water developed in this study.

The extracted DNA is subjected to isothermal amplification via RPA at 37 °C for an optimal reaction time of 20 min. In the presence of target dsDNA containing the *mcyE* gene, the CRISPR-Cas12a system is activated. Cas12a is a class II, type V RNA-guided endonuclease that specifically recognizes and binds to a protospacer adjacent motif (PAM) sequence, 5′-TTTN-3′, located immediately upstream of the target region. Upon successful PAM recognition, the crRNA guides Cas12a to hybridize with the complementary DNA target, resulting in activation of its nuclease function. This activation leads to not only the cis-cleavage of the bound double-stranded target but also a non-specific trans-cleavage activity, wherein Cas12a indiscriminately cleaves nearby ssDNA molecules. This collateral cleavage forms the basis for signal generation in both detection readouts employed in this study. For fluorescence-based detection, an FQ-ssDNA reporter is used. Upon Cas12a activation, the cleavage of this reporter separates the fluorophore from the quencher, generating a fluorescence signal that is proportional to the concentration of the *mcyE* target DNA. This signal can be quantitatively measured by a microplate reader in the laboratory or a portable fluorometer for on-site detection. To complement this quantitative output, an LFA strip was integrated for visual, point-of-use detection. The LFA uses a dual-labeled ssDNA reporter, modified with FAM at the 5′ end and biotin at the 3′ end. In the absence of target DNA, the intact reporter is captured at the control line via biotin-streptavidin interaction, and no signal appears at the test line. In contrast, when the *mcyE* target is present and Cas12a is activated, the reporter is cleaved into two fragments. The FAM-labeled fragment migrates and binds to gold nanoparticle-conjugated anti-FAM antibodies at the test line, producing a visible band, while the uncleaved biotin-labeled fragment (if any left) continues to bind at the control line. Thus, two bands usually indicate a positive result, whereas a single control band signifies a negative result. Overall, this dual-mode RPA-CRISPR/Cas12a detection platform enables both quantitative and qualitative identification of *mcyE* genes in complex lake water matrices, making it highly suitable for on-site or point-of-use monitoring of MC-producing cyanobacteria.

### 3.2. Validation and optimization of the CRISPR/Cas12a crRNA design

Five crRNAs, crRNA 1-5, were designed to target different regions of the *mcyE* gene using the CHOPCHOP tool (http://chopchop.cbu.uib.no/). The designed crRNA sequences are listed in **Table S2**. For each crRNA, the dsDNA template was PCR amplified from genomic DNA extracted from *M. aeruginosa* PCC 7806 with primers designed to amplify ∼1000 bp of the sequence that contains the corresponding crRNA target region. The dsDNA templates were then used for all the crRNA validation tests. The PCR primer designs were validated and optimized through both sequencing and agarose gel electrophoresis. The primer validation and sequences are shown in **Figure S1 and Table S2,** for crRNA 1 as an example. All crRNAs were first evaluated using an in vitro cleavage assay, consisting of the corresponding crRNA, LbaCas12a, and the dsDNA template. The agarose gel electrophoresis (**Figure 2b**) confirmed that all target dsDNAs were cleaved into two distinct fragments within a 30-min incubation, demonstrating that all five crRNAs effectively recognize the target sequences and activate CRISPR/Cas12a cleavage activity. Following this validation, an FQ-ssDNA reporter probe (5′-FAM-TTATT-BHQ1-3′) was introduced into the system to evaluate the trans-cleavage efficiency of crRNAs for optimal sensitivity. As shown in **Figure 2a**, although all five crRNAs triggered fluorescence via trans-cleavage of the nonspecific ssDNA probe, crRNA 1 produced a significantly higher fluorescence signal compared to the others. Therefore, crRNA 1 was selected for CRISPR-Cas12a-based biosensing of the *mcyE* gene. To evaluate the specificity of the constructed CRISPR-Cas12a/crRNA 1 complex, cross-reactivity tests were performed using DNA from closely related genomic regions, including *mcyA*, *mcyD*, and *cpcB*, which are found in both microcystin-producing and non-producing strains of *Microcystis* species. As shown in **Figure 2c**, all these potential interfering targets yielded fluorescence signals comparable to the negative control (i.e., the reporter assay without template DNA), while the *mcyE* gene generated a significantly higher signal. The specificity was also confirmed by agarose gel electrophoresis from the target cleavage tests, which showed no cleavage for any of the potential interfering genes above (**Figure S2**). The completely constructed CRISPR-Cas12a/crRNA complex was illustrated in **Figure 2d**, with the sequences of crRNA 1 and the target region.

**Figure 2.**
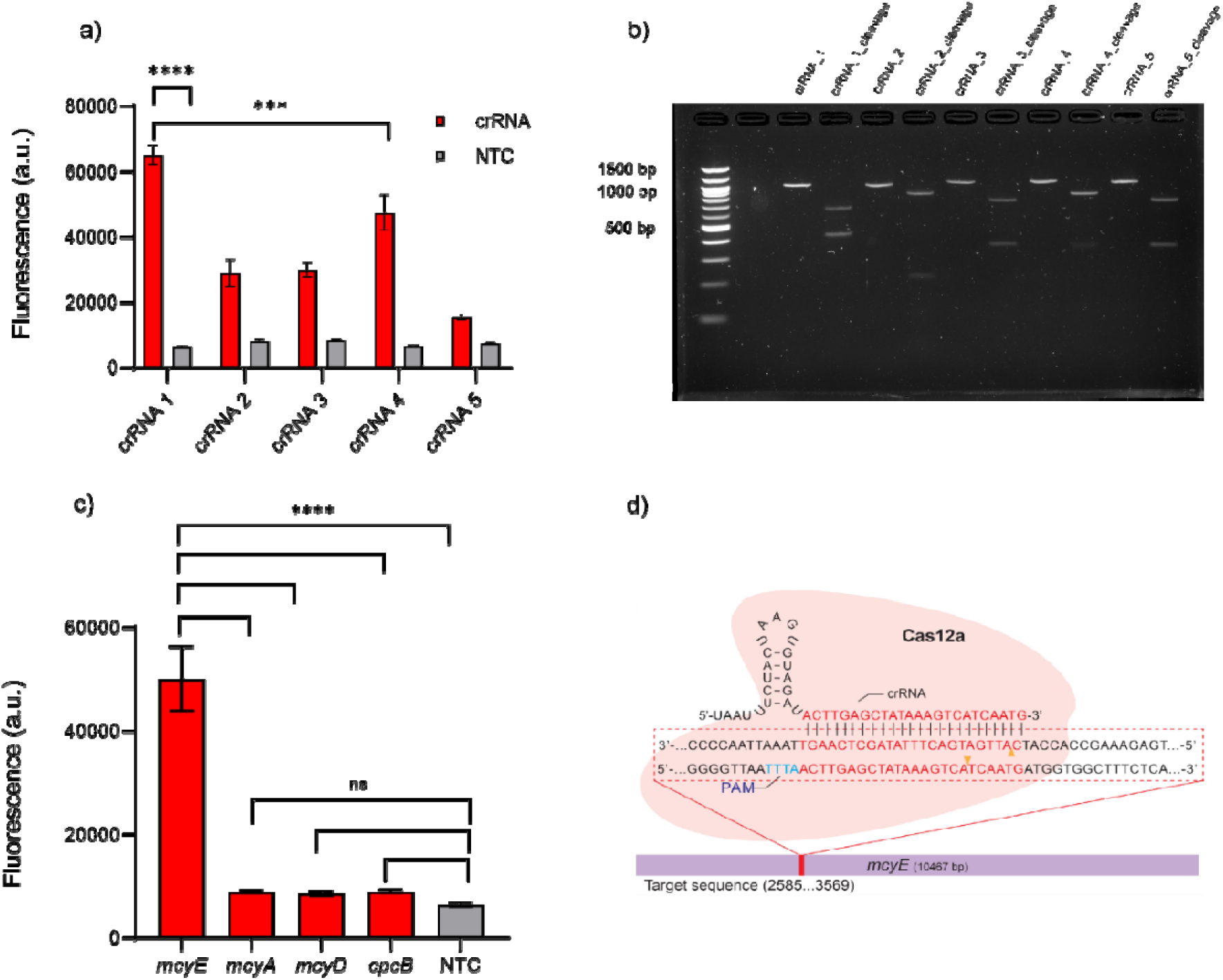
Optimization and validation of the developed method: a) Sensitivity screening for all five designed crRNAs, b) Cleavage activity validation for all five designed crRNAs. The agarose gel electrophoresis image shows the target template dsDNA for each crRNA before and after the cleavage activated by the CRISPR-Cas12a complex. c) Specificity evaluation of the selected CRISPR-Cas12a/crRNA 1 complex by comparing fluorescence (a.u.) of the potential interfering genomic species with the target *mcyE* gene, d) crRNA and target sequence (red) design with the target cleavage sites labeled with orange triangles. Bar graph data represent mean ± SD (n = 3). ****p < 0.0001, ***p < 0.0002, **p < 0.0021, *p < 0.0332, ns > 0.1234 ns = not significant, by one-way ANOVA Tukey’s post-test.

### 3.3. Optimization and analytical performance of the RPA-CRISPR/Cas12a assay

To establish an efficient RPA-CRISPR/Cas12a assay for *mcyE* gene detection, several primer pairs were designed for the RPA preamplification step. Their performance was evaluated by agarose gel electrophoresis and confirmed through sequencing of the amplified products (**Figure S3**). The optimal primer set (**Table S3**) was selected based on its ability to yield a strong, specific amplicon.

Using the selected primers, the RPA incubation time was then optimized with serial dilutions of genomic DNA extracted from *M. aeruginosa* PCC 7806. As shown in **Figure S4**, a 20-min incubation provided the strongest fluorescence signal at lower DNA concentrations, while 15 minutes yielded better performance at higher concentrations. Given that detection sensitivity at low target levels is critical for early-stage HAB monitoring, 20 min was selected as the optimal RPA incubation time. Similarly, the incubation time for the CRISPR/Cas12a-mediated trans-cleavage step was optimized. As shown in **Figure S5**, extending the reaction time from 15 to 30 minutes significantly enhanced fluorescence intensity. Therefore, a 30-min incubation was adopted for the CRISPR/Cas12a reaction to maximize signal generation and assay robustness.

The analytical sensitivity of the RPA-CRISPR/Cas12a assay was evaluated using serial dilutions of extracted genomic DNA ranging from 1.2 to 1.2 × 10^7^ copies/μL. As shown in **Figure 3a**, a concentration-dependent increase in fluorescence signal was observed, with statistically significant differences detected at concentrations ≥1.2 × 10^3^ copies/µL compared to NTC. Although the 1.2 × 10^2^ copies/µL group did not reach statistical significance relative to NTC under one-way ANOVA (Tukey’s post-test), a consistent trend of elevated signal was still observed. The analytical LOD was determined using the 3σ/slope method from the linear calibration curve shown in **Figure 3c**, yielding an LOD of 48.4 copies/µL for extracted genomic DNA. This approach reflects the minimum detectable concentration based on signal-to-noise characteristics and is commonly used in analytical chemistry. The discrepancy between the statistical and calculated LODs may be attributed to signal variation at low target concentrations. Nonetheless, the regression-based LOD provides an estimate of assay sensitivity under optimal conditions and reflects the performance of the assay in controlled measurements. This analytical LOD of the developed RPA-CRISPR/Cas12a assay is also comparable to that of qPCR detection, i.e., 25.7 copies/μL, with the calibration curve shown in **Figure S6**. The specificity of the assay is confirmed in **Figure 3b**. All non-target genes generated comparable fluorescence intensities to the NTC and showed no band at the test line (T line) on LFA strips. Real-time fluorescence monitoring also revealed a clear, concentration-dependent signal increase across the dilution range (**Figure 3d**), confirming the assay’s robust performance over time and dynamic responsiveness. In summary, these results confirm that the developed RPA-CRISPR/Cas12a assay offers a detection sensitivity comparable to qPCR and can reliably detect the *mcyE* gene with high specificity.

**Figure 3.**
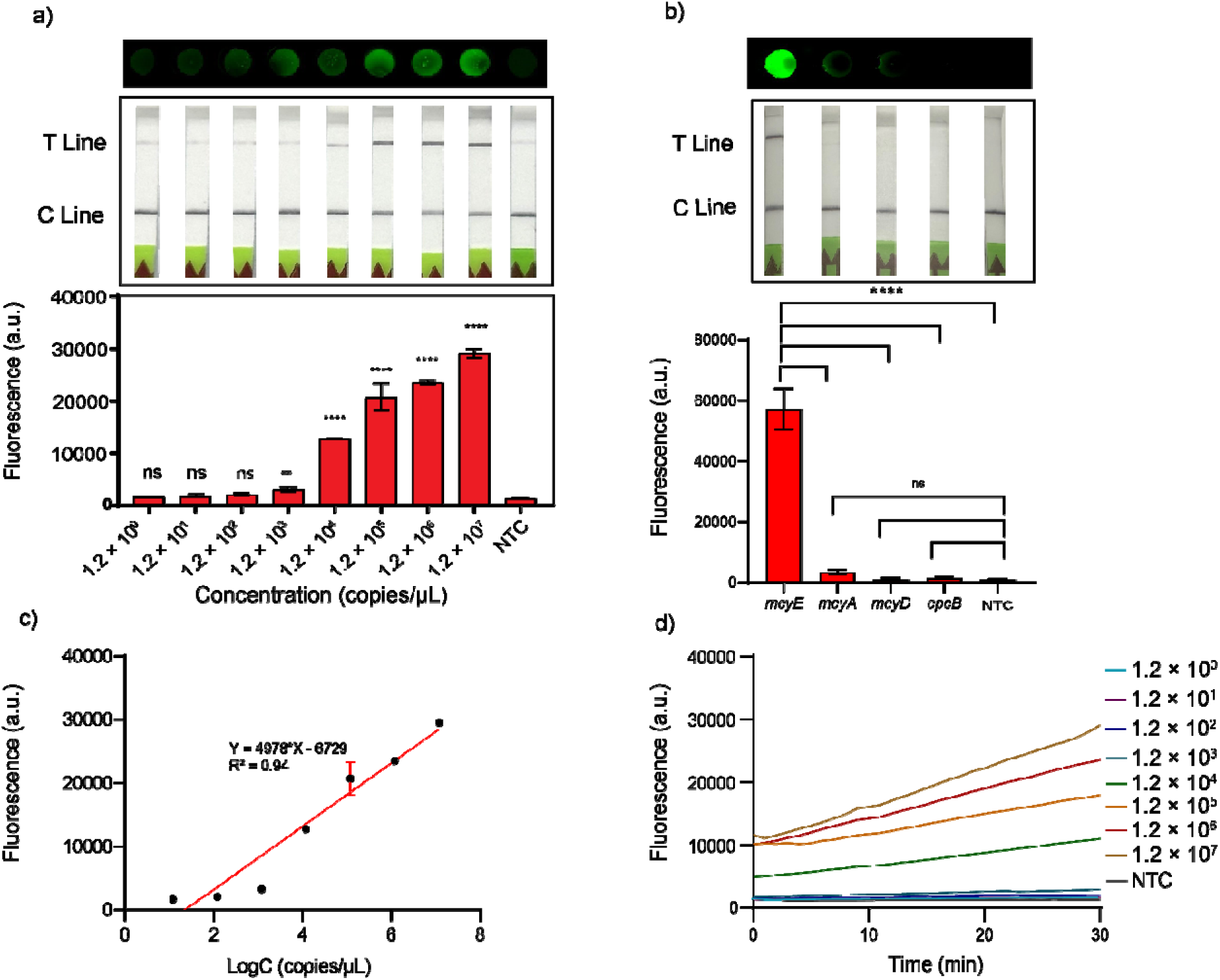
Analytical performance of the developed RPA-CRISPR/Cas12a assay. a) The sensitivity of the RPA-CRISPR/Cas12a biosensing assay for the *mcyE* gene detection from extracted genomic DNA, with fluorescence (top and bottom) and LFA (middle) readout. b) The specificity of the RPA-CRISPR/Cas12a biosensing assay targeting mcyE gene compared with potentially interfering genes, including *mcyA, mcyD,* and *cpcB*. c) The calibration curve of RPA-CRISPR/Cas12a biosensing assay for the *mcyE* gene detection with linear regression equation and correlation coefficient. d) Fluorescence kinetics performance of genomic DNA in the target concentration range of 1.2 - 1.2×10^7^ copies/μL. Bar graph data represent mean ± SD (n = 3). ****p < 0.0001, ***p < 0.0002, **p < 0.0021, *p < 0.0332, ns > 0.1234 ns = not significant, by one-way ANOVA Tukey’s post-test.

### 3.4. Environmental matrix effect on CRISPR/Cas12a-based detection

To evaluate the robustness of the CRISPR/Cas12a biosensing platform under realistic environmental conditions, we systematically investigated the impact of several representative matrix components commonly found in freshwater systems. This included pH, alkalinity (HCO_3_^-^), NOM, nutrient ions (NH_4_^+^ and PO_4_^3+^), and divalent/trivalent metal ions (Ca^2+^ and Fe^3+^), as summarized in **Figure 4a-g**, respectively. The fluorescence intensity generated by Cas12a trans-cleavage activity was used as a direct indicator of assay performance.

**Figure 4.**
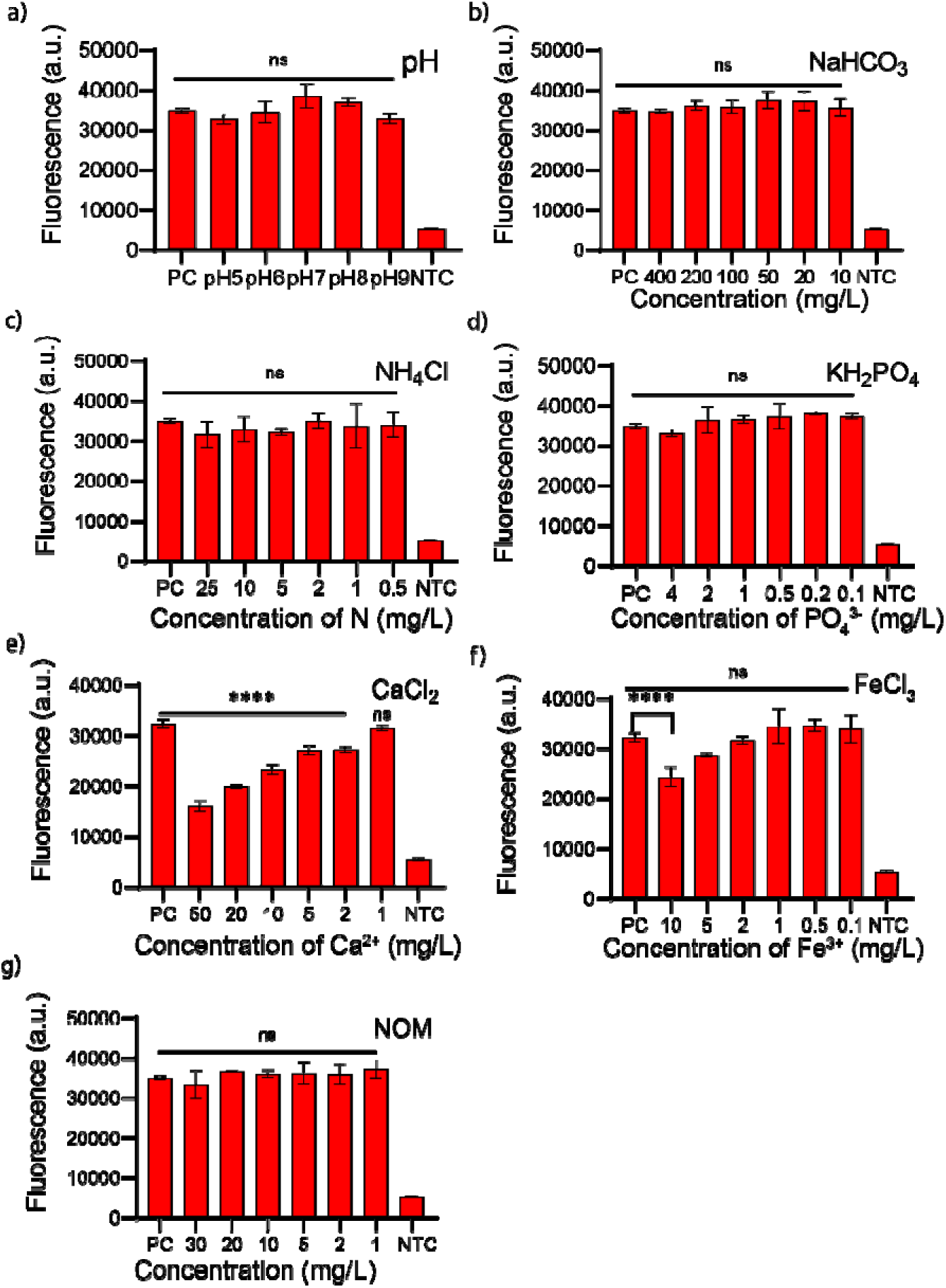
Matrix effects on CRISPR/Cas12a detection, including different matrix factors (a-g) pH, NaHCO_3_, NH_4_Cl, KH_2_PO_4_, CaCl_2_, FeCl_3,_ Natural Organic Matter (NOM). Bar graph data represent mean ± SD (n = 3). ****p < 0.0001, ***p < 0.0002, **p < 0.0021, *p < 0.0332, ns > 0.1234 ns = not significant, by one-way ANOVA Tukey’s post-test.

DNA integrity is known to be pH-dependent, with optimal structural stability maintained between pH 5 and 9. At pH values below 5, protonation of nitrogenous bases leads to depurination and disruption of hydrogen bonding, resulting in strand destabilization. Conversely, at pH values above 9, excess hydroxide ions promote alkaline denaturation through base deprotonation, leading to DNA strand separation and reduced recognition efficiency by crRNA-guided Cas12a complexes. ^25^ These pH-induced structural changes compromise the formation of the R-loop complex essential for Cas12a activation. ^26^ However, the results in **Figure 4a** indicate that within the tested range of pH 5-9, no significant differences in fluorescence output were observed, suggesting stable assay performance under typical environmental pH conditions. Alkalinity, evaluated via NaHCO_3_, also had minimal impact on sensor function (**Figure 4b**). This indicates that moderate buffering capacity in natural water systems does not interfere with the hybridization or enzymatic cleavage efficiency of Cas12a. Similarly, the addition of nutrient ions, ammonium (NH_4_^+^) and phosphate (PO_4_^3-^), did not significantly affect fluorescence output (**Figures 4c and 4d**). Although elevated concentrations of these ions have been reported to influence DNA polymerase activity by destabilizing hydrogen bonds or chelating essential cofactors, such as Mg^2+^, ^27,28^ no measurable inhibition was observed under the tested conditions. These findings suggest that the CRISPR/Cas12a biosensor is tolerant to typical nutrient levels found in eutrophic freshwater systems. In contrast, the presence of divalent and trivalent metal ions demonstrated a clear inhibitory effect. Ca^2+^ and Fe^3+^ added at elevated concentrations significantly suppressed Cas12a activity, as evidenced by a marked reduction in fluorescence intensity (**Figures 4e and 4f**). These ions are known to compete with magnesium for binding at the active site of Cas12a, displacing the essential cofactor and disrupting the catalytic activity of the RuvC domain. ^26^ Furthermore, they may promote DNA aggregation or precipitation, further limiting the accessibility of target sequences to the Cas12a/crRNA complex. These results underscore the importance of controlling divalent and trivalent metal contamination in field applications of CRISPR-based detection. Finally, the influence of NOM was evaluated using environmentally relevant concentrations of NOM standards. While these substances are known inhibitors of DNA-based assays due to their propensity to bind and mask template DNA (Opel et al., 2010), ^29^ the CRISPR/Cas12a platform demonstrated strong resistance to NOM interference. As shown in **Figure 4g**, no statistically significant decline in fluorescence signal was observed across the tested NOM concentrations. This suggests that the biosensing platform remains functional in organic-rich matrices, supporting its practical use in diverse freshwater environments.

It is important to note that this matrix effect investigation was conducted specifically on the CRISPR/Cas12a trans-cleavage step using high concentrations of template dsDNA without any preamplification. However, RPA is also known to be sensitive to several environmental inhibitors. Previous studies have reported that low pH accelerates ATP hydrolysis and disrupts primer binding; high concentrations of phosphate can interfere with primer–template hybridization; and ions such as Ca²⁺ and Fe³⁺ can compete with Mg²⁺ and reduce DNA polymerase activity in RPA reactions. ^30,31^ Additionally, although RPA has demonstrated greater tolerance to NOM than PCR, high NOM concentrations can still hinder amplification efficiency. ^31^

Therefore, to ensure reliable field performance of CRISPR-based biosensors, proper sample pretreatment, including effective cell lysis and DNA purification, is essential. These steps can mitigate the inhibitory effects of environmental matrices on both the preamplification, e.g., RPA, and CRISPR detection phases, improving sensitivity and enabling more consistent results across diverse water sources.

### 3.5. Validation and demonstration of the portable RPA-CRISPR/Cas12a detection platform with lake water

The on-site DNA extraction method was developed to isolate genomic DNA from cyanobacteria in environmental lake water without requiring centrifugation or large laboratory instrumentation (as schematized in **Figure 1b**). To validate this extraction method, its performance was first compared to that of a commercial DNA extraction kit (i.e., QIAamp DNA kit by Qiagen) using both spiked and non-spiked lake water samples, with qPCR as the detection method. The lake water samples were collected from Lake Neatahwanta, Lake Ontario, and Lake Hyde, with cell concentrations shown in **Tables S4**. As shown in **Figure S7a**, lake water samples spiked with *M. aeruginosa* PCC 7806 across a concentration range of 4.7 × 10^3^ to 4.7 × 10^7^ cells/mL yielded comparable C_t_ values between the commercial kit and the developed on-site extraction method. The on-site method achieved a minimum extraction efficiency of 22.6%, demonstrating its ability to recover sufficient DNA for downstream amplification. Furthermore, validation with non-spiked environmental samples from Lake Neatahwanta, Lake Ontario, and Lake Hyde showed that the on-site method could extract DNA with efficiencies of 57%, 52%, and 22%, respectively, relative to the commercial kit (**Figure S7b**). Although the on-site DNA extraction method yielded lower recovery efficiencies compared to a commercial kit, this trade-off reflects its optimization for field use, emphasizing portability, reagent stability, and protocol simplicity over maximal yield. Moreover, the DNA recovered was consistently sufficient for successful RPA-CRISPR/Cas12a detection in both spiked and non-spiked natural water samples, underscoring the method’s functional adequacy for environmental biosensing applications.

Following DNA extraction method validation, the complete RPA-CRISPR/Cas12a detection platform was assembled and tested for a field application format. To assess its sensitivity, the system was evaluated using lake water samples spiked with *M. aeruginosa* over the same concentration range as above for DNA extraction method validation. As shown in **Figure 5b**, a strong fluorescence signal was observed in a dose-dependent manner, with a method LOD of approximately 8.4 × 10^2^ cells/mL, which is much lower than the threshold cyanobacterial concentration of 2 × 10^4^ cells/mL, suggested by WHO for a low probability of adverse health effects from recreational water exposure. ^32^ In addition, the system was applied directly to non-spiked lake water samples and successfully detected the presence of the *mcyE* gene with clear signal readouts (**Figure 5c**). To support true point-of-use or on-site applications, all essential components were integrated into a compact, battery-powered suitcase system (**Figure 5a**). This included reagents for RPA and CRISPR-Cas12a reactions, pipettes, pipette tips, magnets, LFA strips, a thermal block, PES membranes, syringes, and a Qubit 4 portable fluorometer. The built-in rechargeable battery powers the thermal block for up to three hours, and the entire system is car-adapter compatible, enabling flexible field operations. Finally, the fluorescence signal generated from the CRISPR/Cas12a trans-cleavage reaction was successfully quantified using the portable Qubit 4 fluorometer. A standard calibration curve generated with known concentrations of target DNA showed excellent linearity (R^2^=0.9116) as presented in **Figure S8**, demonstrating the fluorometer’s suitability for quantitative on-site readouts.

**Figure 5.**
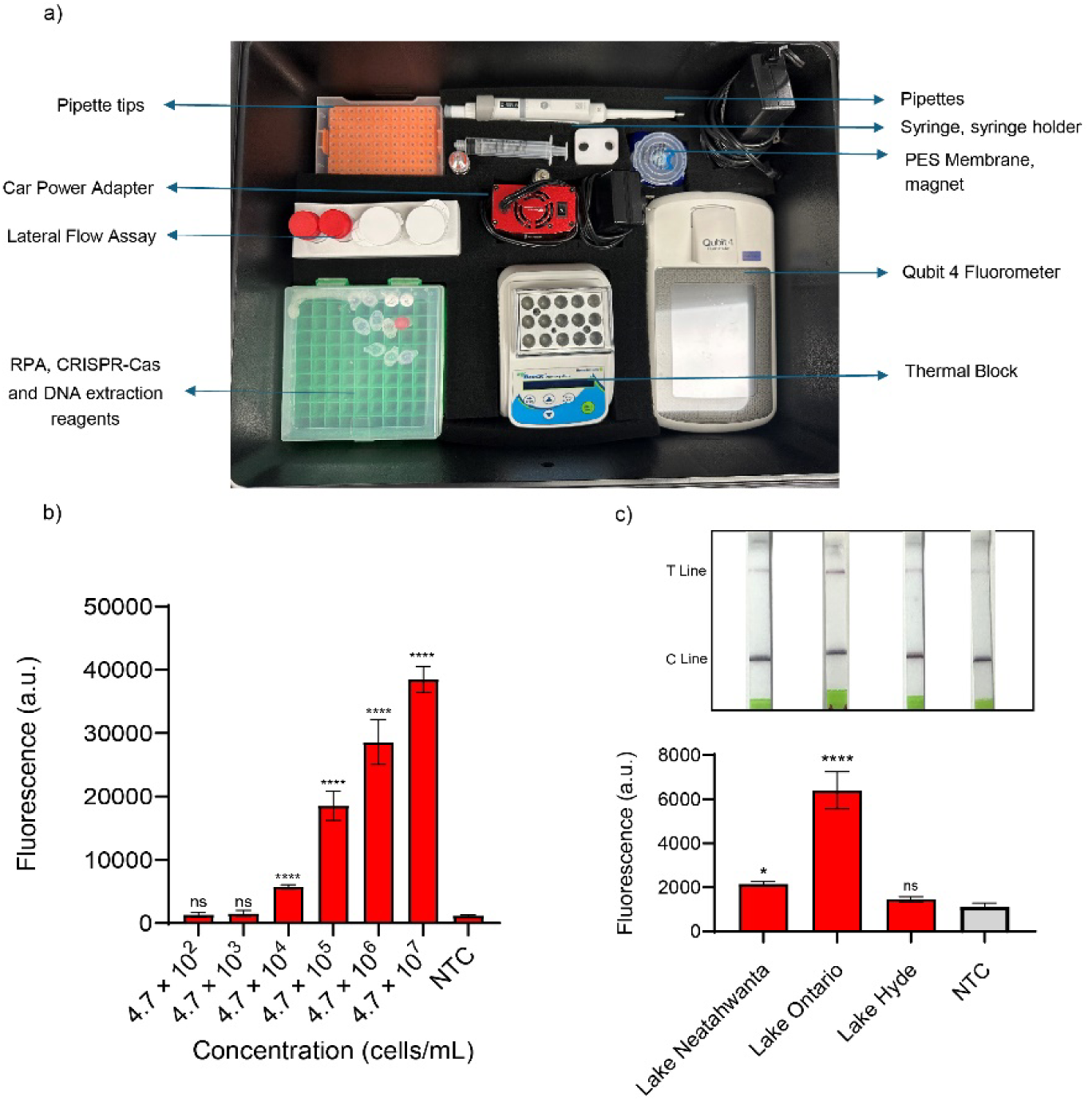
Detection of the *mcyE* gene using the developed biosensing platform. a) Image of the developed suitcase for the on-site detection of the *mcyE* gene in environmental water. b) Serially diluted Microcystis-spiked lake water samples ranging from 4.7 × 10^2^ to 4.7 × 10^7^ cells/mL, c) Real Lake water samples. Bar graph data represent mean ± SD (n = 3). ****p < 0.0001, ***p < 0.0002, **p < 0.0021, *p < 0.0332, ns > 0.1234 ns = not significant, by one-way ANOVA Tukey’s post-test.

## 4. Conclusion

In summary, we developed a complete, field-deployable RPA-CRISPR/Cas12a-based biosensing platform for the rapid and specific detection of the *mcyE* gene from *M. aeruginosa* PCC 7806 in environmental water samples. This integrated system includes a centrifuge-free, simplified on-site DNA extraction method capable of recovering amplifiable genomic DNA within one hour. The DNA yield, although moderate compared to commercial kits, was sufficient to support reliable detection, and the streamlined protocol eliminates the need for laboratory infrastructure with shorter operation time. Coupled with the RPA/CRISPR-Cas12a assay, the platform demonstrated high analytical sensitivity (LOD of 48.4 copies/µL) and specificity, with a total detection time of 50 min, significantly faster than conventional qPCR (∼1.5 hours) while delivering comparable performance. To ensure applicability in complex environmental matrices, we conducted a comprehensive matrix effect evaluation, including pH, alkalinity, NOM, nutrient ions (ammonium and phosphate), and interfering metal ions (Ca^2+^and Fe^3+^). To our knowledge, this is the first systematic investigation of environmental influences on CRISPR-based sensing for cyanotoxin gene targets. Given that CRISPR diagnostics in environmental monitoring are still in their early stages, these insights can inform future biosensor design, assay optimization, and deployment strategies for a wide range of environmental targets. Importantly, this work underscores that the real-world applicability of biosensors, particularly for biological targets in environmental monitoring, depends not only on analytical sensitivity but on the integration of all workflow components, from sample pretreatment to signal readout. Developing the sensing chemistry alone is insufficient; without robust upstream DNA extraction and downstream detection mechanisms compatible with field conditions, practical deployment remains unattainable. The portable suitcase system developed in this study consolidates all necessary instruments and reagents into a single operational unit, making true point-of-use detection possible.

For future development, continued innovation is needed to simplify the platform further. This includes the development of more efficient and autonomous on-site sample pretreatment technologies and the advancement of preamplification-free CRISPR detection strategies to reduce assay complexity and accelerate time-to-result. Together, these efforts will enhance the feasibility and scalability of CRISPR-based biosensors for real-time environmental monitoring and early warning systems targeting harmful algal blooms and other biological contaminants.

**There are no conflicts of interest to declare.**

## Acknowledgements

This project was supported by the New York State Center of Excellence in Healthy Water Solutions [C190175] and the National Science Foundation [23472070]. We thank Anna Malos for his assistance in preparing samples for the investigation of environmental matrix effects on the CRISPR/Cas12a-based biosensing technique. We also thank Dr. Yaqi You, Hanyue Yang, Amina Furrkurkh Mughal, and Athkia Fariha for their help on sampling at Lake Neatahwanta, NY.

## Appendix A

Supporting Information.

